# Role of gene body methylation in coral acclimatization and adaptation

**DOI:** 10.1101/184457

**Authors:** Groves Dixon, Yi Liao, Line K. Bay, Mikhail V. Matz

## Abstract

Gene body methylation (GBM) has been hypothesized to modulate responses to environmental change, including transgenerational plasticity, but the evidence thus far has been lacking. Here we show that coral fragments reciprocally transplanted between two distant reefs respond with genome-wide increase or decrease in GBM disparity among genes. Surprisingly, this simple genome-wide adjustment predicted broad-scale gene expression changes and fragments’ fitness in the new environment. This supports GBM’s role in acclimatization, which may consist in modulating the expression balance between environmentally-responsive and housekeeping genes. At the same time, constitutive differences in GBM between populations did not align with plastic GBM changes upon transplantation and were mostly observed among *F*_ST_ outliers, indicating that they arose through genetic divergence rather than through transgenerational inheritance of acquired GBM states.

**One-sentence summary:** Genome-wide shifts in gene body methylation predict gene expression and fitness during acclimatization but do not contribute to epigenetic divergence between populations.

---

GBM is a taxonomically widespread epigenetic modification the function of which remains unclear (*1, 2*). GBM is bimodally distributed among genes: it is high in ubiquitously expressed housekeeping genes and low in inducible genes (*2, 3*). Only the detrimental effect of GBM is well understood: GBM causes hypermutability in protein-coding regions (*4*). Indeed, in humans, GBM is the primary driver of deleterious parent-age-related mutations (*5*). To merit pervasive evolutionary conservation, the fundamental biological function of GBM must be important enough to outweigh this risk (*1*). Among putative cellular functions of GBM, suppression of intragenic transcription initiation is currently the most studied (*6, 7*) but still remains controversial (*8*). The ecological genomic literature has long expected that GBM (as well as other epigenetic marks) respond to the environment and assist acclimatization by modulating gene expression, possibly across generations (*9–11*). Still, thus far the only well-documented case of GBM responding to the environment and resulting in phenotypic change is caste determination in social insects depending on diet (*12*). Beyond this highly taxon-specific example, GBM response to the environment has been reported in a recent paper on coral acclimatization to acidic conditions (*13*); notably, GBM change did not specifically affect genes expected to be involved in such acclimatization. One way to address the role of GBM in adaptation would be to examine differences between populations, as has been recently done in plants (*14, 15*); however, this interpretation requires disentangling the effect of environment from the effect of genetic divergence among populations (*16*). As for transgenerational plasticity, for GBM to be involved in this process it must be both responsive to the environment and heritable. This somewhat contradictory combination of properties has not yet been demonstrated for GBM in any study system.

Here, we used a reciprocal transplantation framework (*17*) to test for the roles of GBM, gene expression, and genetics in local adaptation and acclimatization in a reef-building coral *Acropora millepora*. In corals, the possibility of clonal replication makes disentangling the effects of genotype and environment very straightforward, solving a major problem of ecological epigenetics (*16*). We compared corals from two reefs, Orpheus Island in the central sector of the Great Barrier Reef (GBR) and Keppel Island in the south of GBR (Fig. 1 A). These reefs are notably different in temperature (Fig. 1 B) as well as several other abiotic parameters (*18*) and host slightly genetically divergent *A. millepora* populations (*19, 20*). Fifteen coral colonies from each reef were halved and reciprocally transplanted between the sites for three winter months (Fig. 1 A, B). In this way, each individual genotype was simultaneously exposed to two distinct reef conditions. After three months in the field, tissue samples were collected from each fragment and assayed for genome-wide gene expression using TagSeq (*21*) and for DNA methylation using MBD-seq (*22*). These data were analyzed in the context of several fitness proxies and genetic distances between individuals (based on genetic polymorphisms detected in the MBD-seq data). All experimental corals were dominated by *Symbiodinium* clade C, which was established by analyzing Symbiodinium-matching MBD-seq reads.

**Figure 1.**
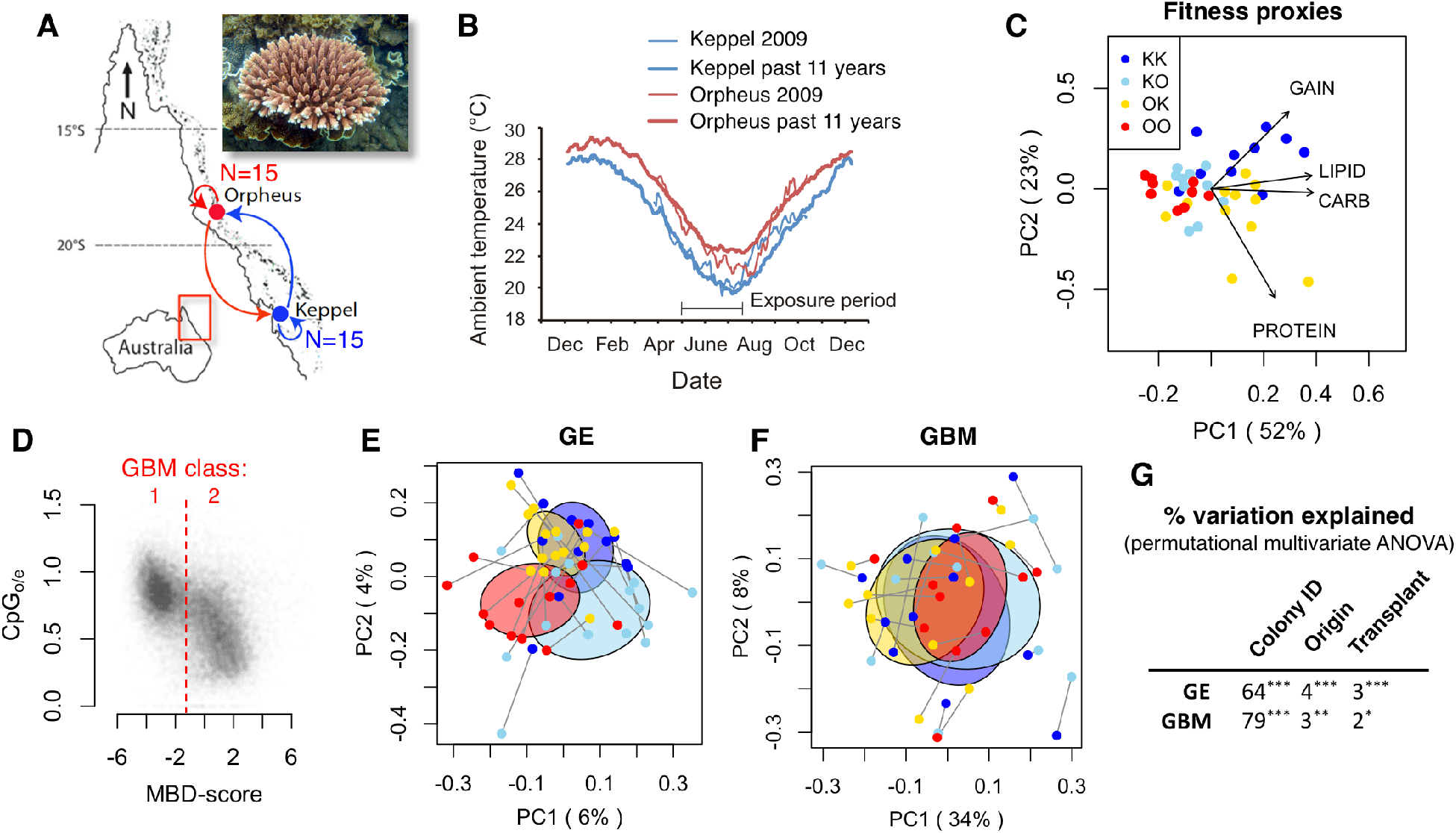
Overview of the experiment. A: Design of reciprocal transplantation. Inset – *Acropora millepora*. B: Temperature profiles at the transplantation sites, historical (thick lines) and during the year of experiment (thin lines). C: Principal component analysis of the four fitness proxies: weight gain, lipid, protein and carbohydrate content. In the sample names, the first letter is the origin location and the second letter is transplantation location. D: Scatter plot of the measure of depletion of CpG dinucleotides (ratio of observed vs. expected number of CpGs, CpG_O/E_) against MBD-score (log_2_ of fold-difference in read counts between MBD-captured and flow-through fraction) for each gene. The MBD-score threshold defining the two GBM classes is indicated. E and F: Principal coordinate analysis of gene expression (E) and gene body methylation (F). Lines connect fragments of the same original colony; the color scheme follows the legend on panel C. For GE, PC1 corresponds to the colony’s origin and PC2 - to the transplantation site; no such partitioning is visible for GBM. G: Percentage of variation in GE and GBM explained by colony identity (i.e., by similarity among clonal fragments, or broad-sense heritability), colony’s origin and transplantation site. *** *p* < 0.001; ** *p* < 0.01; * *p* < 0.05.

For both Keppel- and Orpheus-origin corals fitness proxies were higher at Keppel, possibly due to higher concentration of inorganic nutrients there (*18, 19*). For Keppel-origin corals the difference in performance was predominantly in weight gain (i.e., skeletal growth) while for Orpheus-origin corals the difference was mostly in internal stores (lipids, proteins and carbohydrates, Fig. 1 C). This indicates that (i) Keppel and Orpheus environments are indeed different from corals’ perspective, (ii) both populations are capable of plastic responses to these differences, and (iii) there is also constitutive divergence in physiology between populations.

To determine the characteristic level of GBM for each gene we used MBD-seq results for 12 samples (three per each experimental group, KK, KO, OK, and OO) for which both MBD-captured and flow-through fraction were sequenced. Logarithm with the base 2 of each gene’s abundance in captured vs. flow-through fractions (“MBD-score” (*22*)) exhibited the expected bimodal distribution (Fig. S1 A) and showed strong correlation (Fig. 1 D) with the established proxy of historical methylation level, the ratio of observed to expected numbers of CpG dinucleotides (“CpG_O/E_” (*3*)). In addition to this genome-wide validation of our methylation data, we validated GBM results by bisulfite-sequencing 13 amplicons representing different MBD-score classes (Fig. S1 B, C). Genes were assigned to either low-methylated class (class 1) or high-methylated class (class 2) based on the MBD-score threshold indicated on Fig. 1 D. Relative quantification results based only on MBD-captured read data were nearly identical to those obtained when using both captured and flow-through fractions (Fig. S2). For the remaining 32 samples, we sequenced only the MBD-captured fraction.

Multivariate analysis of gene expression (GE, Fig. 1 E) and GBM (Fig. 1 F) revealed that GBM is more consistent across fragments of the same original colony than GE, resulting in the estimate of broad-sense heritability of 0.79 compared to 0.64 for GE (Fig. 1G, Fig. S3 and S4). In addition, significant effects of origin and transplantation site were observed for GE and GBM, both more pronounced for GE (Fig. 1 E and G).

Surprisingly, the change in GBM in response to transplantation from Orpheus to Keppel consisted mainly in genome-wide reduction of disparity between high- and low-methylated genes: highly-methylated genes became less methylated and low methylated became more methylated (Fig. 2 A). This change was mirrored by less pronounced but clearly reciprocal GBM adjustment in Keppel corals transplanted to Orpheus (Fig. 2 B, D) and recapitulated the difference between fragments planted in their native environment (Fig. 2 C). Both exons and introns underwent this change (Fig. S5). Absolute changes in methylation included both positive and negative shifts (Fig. S6 A, B), indicating that the change in relative GBM levels among genes is not due to biased genome-wide methylation increase or decrease. Methylation levels of intergenic regions and repeated elements remained relatively constant compared to genic regions (Fig. S6 C, D).

**Figure 2.**
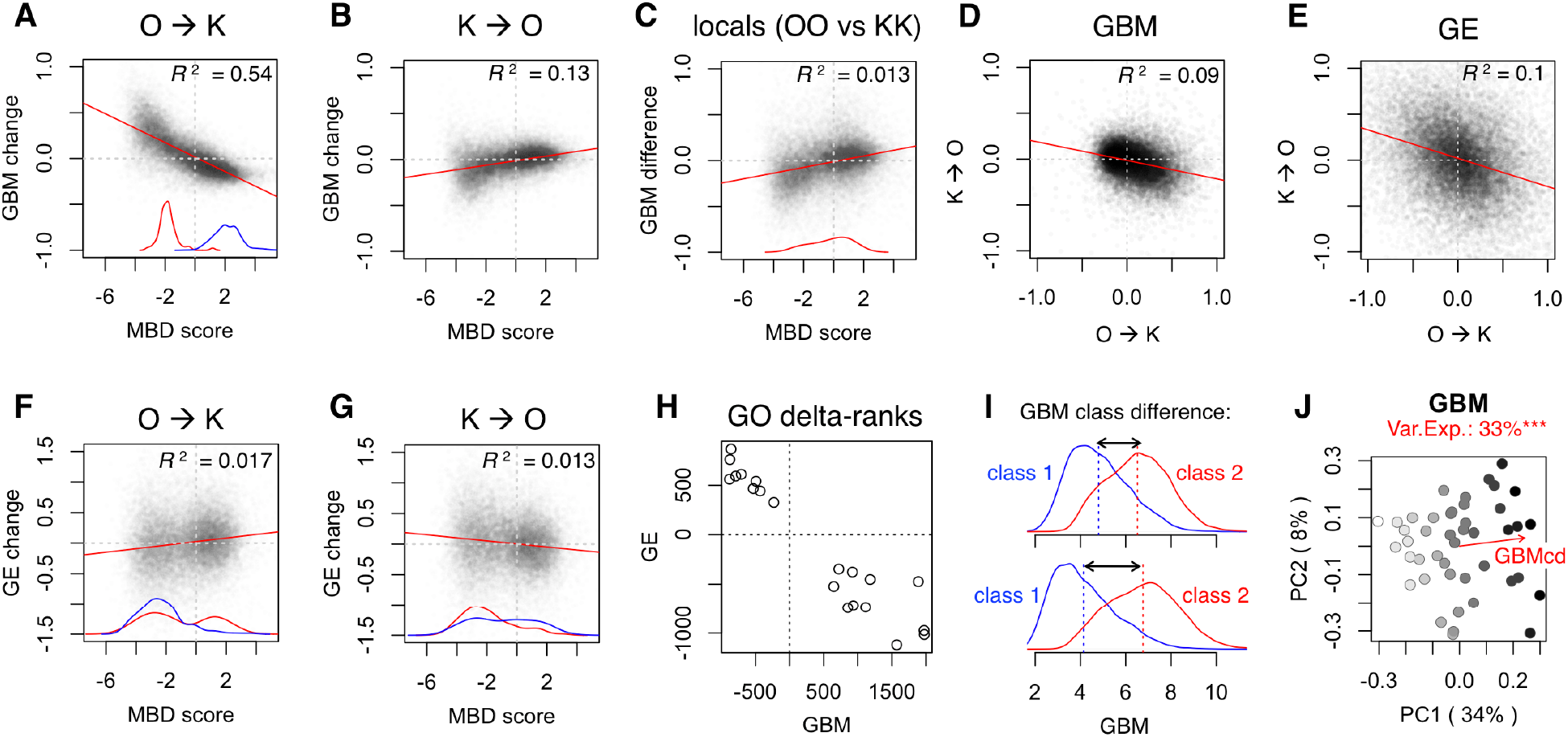
Changes in gene body methylation (GBM) and gene expression (GE) in response to transplantation (“away” relative to “home”). On all panels, GE and GBM measures are log_2_ fold-changes. A though C: Differences in GBM in fragments transplanted from Orpheus to Keppel (A), from Keppel to Orpheus (B), and fragments planted in their home environment (C) plotted against MBD-score for each gene. Red and blue curves near the x-axis are density plots of significantly up- and down-regulated genes (there were none for the K➔O experiment). D and E: Correlation between GBM (D) and GE (E) changes in reciprocally transplanted fragments. F and G: Change in GE in fragments transplanted from Orpheus to Keppel (F) and from Keppel to Orpheus (G) plotted against MBD-score for each gene (compare to panels A and B). H: Plot of delta-ranks for 20 Gene Ontology categories that were significantly enriched with either up-regulated genes (positive delta-rank) or down-regulated genes (negative delta-rank) in both GE and GBM analyses. I: Calculation of GBM class difference (GBMcd), a proxy of genome-wide GBM disparity among genes. Each panel shows two density plots of GBM levels for the two GBM gene classes (see Fig. 1 D), with the mean for each class indicated by a dashed line. The top panel shows the sample with the lowest GBMcd, bottom panel – the sample with the highest GBMcd. J: Principal coordinate biplot (the same ordination as on Fig. 1 F) colored according to GBMcd. The purple vector shows the direction of correlation of GBMcd with the ordination. All correlations with reported R^2^ are significant at the p<0.0001 level.

GE response was also reciprocal between transplantation directions but demonstrated much broader range of fold-changes compared to GBM response (Fig. 2 D). Remarkably, gene expression changes correlated with low-and high-methylated gene classes and were in the opposite direction relative to GBM changes (Fig. 2 F, G). Although at the gene level the correlation between GE and GBM was weak (but still highly significant, Fig. S7), it was very strong when comparing broader functional groups of genes (Gene Ontology categories, Fig. 2 H and Fig. S8).

To characterize GBM disparity among genes in each sample, we computed GBM class difference (“GBMcd”) as difference in mean log_2_(GBM) between high- and low-methylated gene classes (Fig. 2 I). This measure aligned nearly perfectly with the first principal component of GBM variation for the whole experiment, explaining 33% of total variation (Fig. 1 F and Fig. 2 J, which show the same PCA ordination but are colored differently). For comparison, the next principal component explains only 8% of GBM variation.

Our data do not support transgenerational inheritance of acquired GBM changes. If plastic changes in GBM were transmitted across generations, one would expect constitutive GBM differences between Orpheus- and Keppel-origin corals to parallel the GBM response induced upon transplantation. Instead, constitutive differences between Orpheus and Keppel (differences between all Orpheus-origin and all Keppel-origin fragments, irrespective of transplantation site) were in the opposite direction compared to plastic changes (compare Fig. 3 A and Fig. 2 C), did not predominantly affect genes at the opposite ends of MBD-score spectrum (compare density curves on Fig. 2 A and Fig. 3 A), and were positively rather than negatively associated with differences in GE (Fig. 3 B). In addition, genes showing significant constitutive differences in GBM also showed elevated *F*_ST_ (Fig. 3 C) unlike genes undergoing significant plastic changes (Fig. S9). This indicates that the mechanism giving rise to constitutive GBM changes between populations is unrelated to the mechanism of plasticity and is most likely genetic divergence.

**Figure 3.**
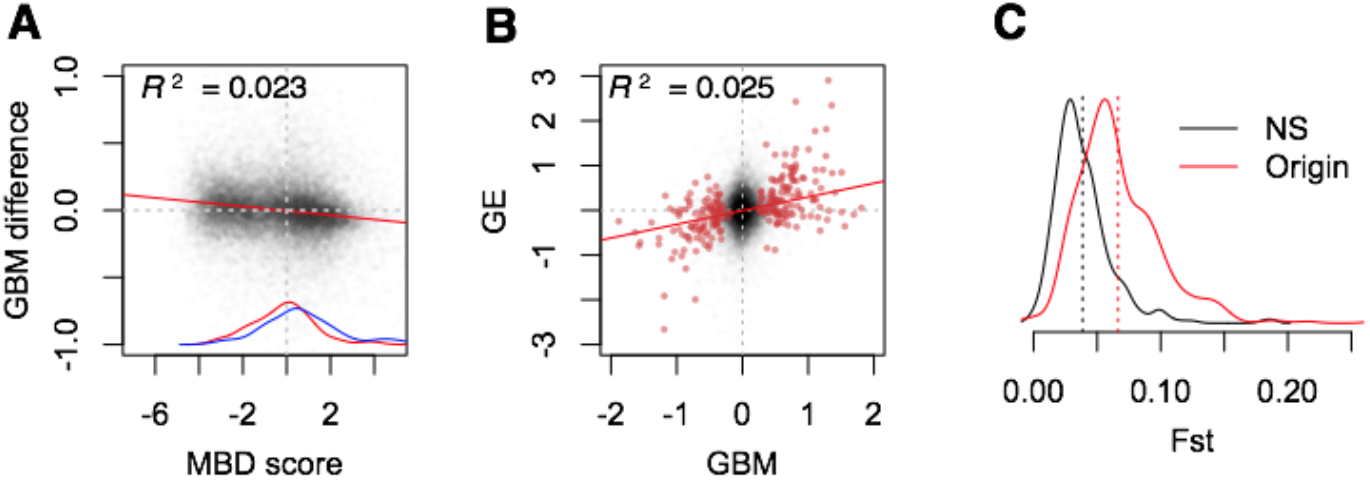
Constitutive log_2_ fold-differences in gene body methylation (GBM) and gene expression (GE) between populations (Orpheus relative to Keppel). A: Difference in GBM plotted against MBD-score for each gene. Red and blue curves by the x-axis are density plots of significantly up- and down-regulated genes. B: Correlation between GBM and GE differences. Red points are genes significantly different in GBM at 10% FDR level. This correlation is positive, while correlations between plastic GBM and GE changes are negative (Fig. 2 H and Fig. S7). E: Density plot of between-population *F*_ST_ for genes showing significant constitutive difference in GBM (red line) compared to 500 randomly chosen non-significant genes (black line). Dashed vertical lines mark the mean of each dataset. All correlations with reported R^2^ are significant at p<0.0001 level.

To see whether plastic GBM changes were related to acclimatization (rather than, for example, overall stress in response to transplantation) we used Differential Analysis of Principal Components (DAPC, Fig. 4 A-C) to test whether GBM similarity to the local population (Fig. 4 D) predicted the transplanted fragment’s fitness in the new environment. The resulting “GBM similarity to locals” was highly correlated with the first principal component of measured fitness proxies, the most strongly with the weight gain (Fig. 4 E). This correlation held both before and after (as shown on Fig 4 E) controlling for overall higher fitness of Orpheus corals transplanted to Keppel. This result indicates that a coral’s GBM profile, despite being highly variable among individuals (Fig 1 F, G), also reflects adaptation and acclimatization to local environmental conditions. Interestingly, the same analysis performed for GE and genotypes (GT) revealed no relationships with fitness (Fig. 4 E). One explanation of this could be the difference in time scales on which the effects of GE and GT on fitness would be detectable: compared to the three-month length of our experiment, GE might be linked to fitness on shorter time scales, and GT – on longer time scales.

**Figure 4.**
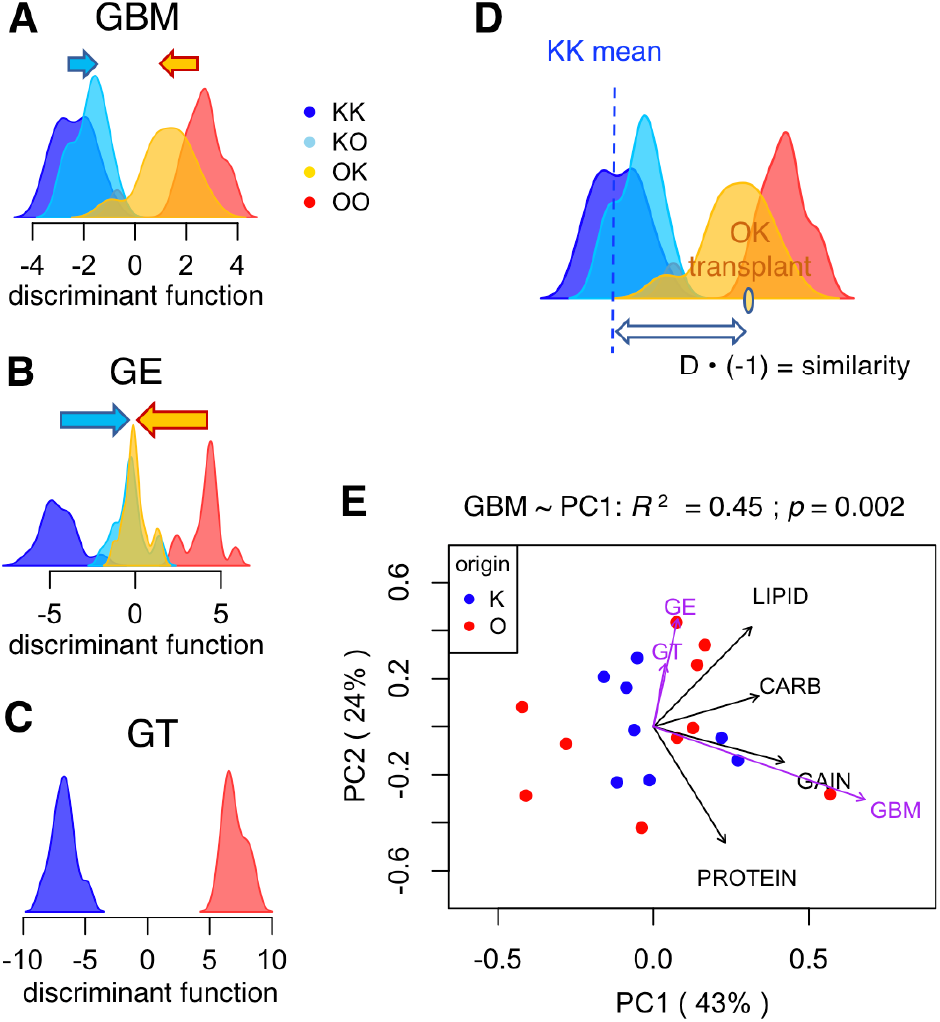
Relationship between gene body methylation (GBM), gene expression (GE), and genotype (GT) with fitness proxies in corals transplanted into novel environment. A-C: Density plots of discriminant function values for each variable. The functions were developed to distinguish between coral fragments in their native environment (KK vs. OO) and applied to transplanted fragments (KO and OK) to quantify their adaptive plasticity (ability to shift towards local values). Arrows indicate mean plasticity. D: Calculation of “similarity to locals” for each transplanted fragment. E: Principal components biplot of fitness proxies in fragments transplanted to another reef, corrected for the overall higher fitness of Orpheus corals transplanted to Keppel. Fit of “similarity to locals” for GBM, GE and GT onto this ordination is shown as purple vectors scaled by correlation coefficient. Only GBM fit is statistically significant.

Association between plastic GBM changes and gene expression changes (Fig. 2 F - H) prompts the question: which one is likely to be the leading cause? The magnitude of GBM changes is much lower than changes in GE (Fig. 2 D, E), which suggests that GE might be the primary driver. However, considering the simplicity of plastic GBM changes - genome-wide exaggeration or mitigation of GBM disparity among genes - it is more parsimonious to assume that this is the leading change driven by the environment, which then translates into broad-scale functional shifts in gene expression (Fig. 2 H and Fig. S8). In this way, genome-wide adjustment of gene activity could be achieved by modulating a single process that governs the disparity in GBM among genes.

Our results suggest that GBM disparity controls the balance between expression of two broad gene classes: the low-methylated environmentally-responsive genes and highly-methylated housekeeping genes. In the higher-quality environment of the Keppel island, where corals attain higher fitness (Fig. 1C), GBM disparity change is associated with suppression of the environmentally responsive genes and up-regulation of housekeeping genes (Fig. 2 F and Fig. S8). Conversely, at the low-quality Orpheus Island the balance shifts the other way, from housekeeping to environmental responsiveness (Fig. 2G and Fig. S8). We propose that mediating such coarse adjustment of physiology in response to the environment is the ecological function of GBM.

## Materials and Methods

### Sample sizes

The experiment started with 15 colonies from each of the two sites, which were halved and transplanted, resulting in a total of 60 fragments distributed across four study groups 15 samples each: two “natives” (KK and OO), and two “transplants” (OK and KO, Fig. 1C). Although not all original samples were successfully analyzed using all approaches (Table S1), in the end each of these four groups included 11 fragments representing unique coral genotypes and paired by genotype between “home” and “away” groups that were analyzed for GBM, gene expression, genotype, and fitness proxies.

### Reciprocal Transplantation Experiment

Field work was conducted with permission from the Great Barrier Reef Marine Park Authority (Research permit G09/29894.1) as described previously (*23*). Reciprocal transplantations were undertaken between two environmentally distinct study sites (Miall Island in the Keppel Island group: 23°09S 150°54E and Hazard Bay on Orpheus Island 18°37S 146°29E) separated by 4.5 degrees of latitude on the Great Barrier Reef (Figure 1A). On the 23rd of April (Orpheus) and 4th of May (Keppels) 2010 fifteen colonies were collected from wild populations from each site and split in two. One half of each colony was replaced in its native habitat, while the second half was transplanted to the alternate study site. Samples from all coral fragments were collected at midday after three months (9th July 2010 at Orpheus, 14th July at Keppels) frozen in liquid nitrogen, then transferred into RNAlater (Ambion, Austin, TX, USA) for gene expression and DNA methylation profiling.

### Coral growth rate

Total colony skeletal growth was evaluated using wet buoyant weight as in Jokiel, Maragos, & Franzisket (1978). Measurements from beginning and end of the experiment were expressed as percent daily weight gain.

### Coral condition

The energetic condition of corals was determined by the analysis of total protein, carbohydrate and lipid content. Coral tissue was removed from an average of 8.6 cm^2^ of coral branch (mean + SE = 0.18) using an air gun in 12 – 15 ml of 0.2 μM filtered sea-water (FSW). The coral slurry was homogenized and centrifuged at 1500 g for 5 min at 4°C to separate coral and endosymbiotic algal components. The coral slurry was aliquoted for protein, carbohydrate and lipid assays and frozen at −20°C (10 ml for lipid, 0.5 ml for protein and carbohydrate). The algal component was re-suspended in 2.75 ml of 0.2 μM FSW. Coral tissue surface area was determined from cleaned (10% bleach for 24 hrs) skeletons of water-blasted samples using a twice dip paraffin wax method (*25*). Surface area of coral branches was determined by the weight gain of the second wax layer from a session specific standard curve (R^2^ were 0.97 – 0.98) using seven nylon cylinders of known surface area (5 – 48 cm^2^).

### Protein analysis

Total protein was extracted from a 0.5 ml aliquot using 0.5 ml 1M NaOH. Samples were incubated at 90°C for one hour and spun at 3000 g for 5 minutes to separate cell-debris from the solution and 500 μL clear supernatant was transferred onto a 96-well format. Protein concentration was quantified in three technical replicates of 50 μl of coral protein extract using a microplate Peterson – Lowry assay following the manufacturer’s recommendations (Sigma: TP0300). Blank 0.5 M NaOH samples and BSA protein standards (50, 100, 200, 300 and 400 μg / ml) were run in duplicates on each plate and absorbance was read at 595 nm on a spectrophotometer. Standard curves had R^2^ of 0.97 – 0.99. The coefficient of variation (CV) was calculated among technical replicates and deemed acceptable when <10%. For samples with a CV > 10%, single deviant outliers (>1 SD deviant from other two measurements) were identified and omitted. Total protein content per sample was expressed as average content (mg) of the technical replicates per cm^2^ of coral branch surface area.

### Carbohydrate analysis

Total carbohydrate content estimates were obtained from the average of three technical replicates of 50 μL coral slurry using D-glucose as a standard (*26*). Blank water samples and BSA protein standards (20, 50, 100, 200, 500 and 1000 μg/ml) were run in triplicate on each plate and absorbance was read at 485 nm on a spectrophotometer. Standard curves had R^2^ of 0.96 – 0.99 and total carbohydrate content per sample was expressed in average mg of the technical replicates per cm^2^ of coral branch surface area after quality control as described for proteins.

### Lipid analysis

Lipids were extracted using a modified version of Harland et al. (1993). Ten ml of coral slurry was freeze dried overnight then homogenised by vortexing with 2 × 5 mL of 2:1 dichloromethane:methanol (DCM:MeOH), each left for 24 hours at 4°C. Extracts were filtered to remove any debris using a glass filter (Whatman GF/C) and purified once with 5 ml 0.88% KCl:H_2_O and thrice with 1:1 MeOH:H_2_O. Total lipid content was determined gravimetrically from dried samples (60°C overnight) in pre-weighed acetone washed aluminum trays. Total lipid content was expressed in mg per cm^2^ of coral branch surface area.

### MBD-seq library preparation

DNA was isolated from adult holobiont tissue using dispersion buffer (4 M guanadine thiocyanate, 30 mM sodium citrate, 30 mM β-mercaptoethanol) followed by phenol chloroform purification and a final cleanup with Zymo Genomic DNA Clean and Concentrator-10 kit (Catalog No D4011). Genomic DNA was sheared using a Misonix Sonicator 3000 to a size range of ~200 to 800 bp checked by gel electrophoresis. Enrichment reactions were performed using the MethylCap kit (Diagenode Cat. No. C02020010) with an initial input of 2 μg of sheared DNA per reaction. The methylated fraction was eluted from the capture beads in a single step using High Elution Buffer. Library preparation using NEBNext^®^ DNA library Prep Master Mix Set (E6040L) and sequencing on an Illumina HiSeq4000 was performed at the University of Texas Genome Sequencing and Analysis Facility. Our total sample size was N=44 (22 colonies divided in half giving 11 samples per treatment group). For the majority of these we sequenced only the enriched library eluted from the capture beads. Fold coverages from these captured libraries were used to estimate relative differences in GBM between samples. For a subset of 12 of the 44 samples, we sequenced both the captured and flow-through fractions. Fold differences between these captured and flow-through libraries were used to estimate absolute levels of methylation across genes. We did this for only a subset of samples because we were primarily interested in relative differences in GBM between groups. As relative differences could be reasonably well assessed without sequencing the flow-through (Figure S2), we chose to focus our sequencing resources on increasing sample size rather than more thorough estimates of absolute methylation levels.

### Repeated elements annotation

*De novo* identification of repeated elements in *A. digitifera* genome was performed using RepeatModeler (http://www.repeatmasker.org/RepeatModeler/, version 1.0.10). RepeatModeler launches two repeat finding programs, RECON 1.05 (*28*) (http://eddylab.org/software/recon/) and RepeatScout 1.0.5 (*29*) (https://bix.ucsd.edu/repeatscout/), to generate a genomic database and upon which to build, refine and classify consensus models of putative interspersed repeats. This pipeline produced a total of 2,858 non-redundant repeat families in recent *A. digitifera* genome sequence. These sequences were then used as the custom library to search the *A. digitifera* genome using RepeatMasker (http://www.repeatmasker.org/). Approximate 7.33% and 5.47% of the genome was annotated to be composed of retrotransposon and DNA transposon elements, respectively.

### MBD-seq data processing

Forty four MBD-seq libraries were prepared from reciprocally transplanted coral fragments. Sequencing produced 980 million raw reads with a mean of 16 ± 0.65 SEM million reads per sample. Raw reads were trimmed of non-template sequence using Cutadapt (*30*) and quality filtered using Fastx toolkit (http://cancan.cshl.edu/labmembers/gordon/fastx_toolkit/). Adapter trimming and quality filtering reduced these the total read count 940 million, mean = 15 ± 0.64 SEM million reads per sample. The reference genome and annotations (version 1.1) for *Acropora digitifera* (*31*) were downloaded from NCBI. Trimmed and filtered reads were mapped to this concatenated reference using Bowtie2 (*32*). To ensure that using a reference from an alternate species did not severely impair mapping, we compared the mean mapping efficiency against the *A. digitifera* reference with that of a draft genome sequence for *A. millepora* produced by David Miller and coworkers (James Cook University). Mean mapping efficiency against the *A. digitifera* (78.4 ± 0.7% SEM) reference was only 5.2% lower than that of *A. millepora* (83.6 ± 0.8% SEM), hence sequence differences between the two species did not appear to substantially impair read mapping. Following mapping, PCR duplicates were removed using Picard (https://broadinstitute.github.io/picard/). Mean duplication frequency was 14.3 ± 0.5% SEM. The BAM files were filtered to retain only highly uniquely aligned reads (mapping quality 30, or 0.1% chance of non-unique alignment) using samtools (*33*), after which the number of reads overlapping various genomic regions - exon, intron, genic (exon + intron), repeated elements longer than 500b, and repeat-free intergenic regions at least 1 kb long and at least 2 kb away from any gene - were counted using BEDtools (*34*).

### MBD-score and GBM gene classes

For 12 samples (3 per each experimental group) we quantified absolute levels of GBM as the log_2_ fold difference in coverage between captured and flow-through DNA fractions while controlling for genotype, as described in Dixon et al. (2016). These values were used as gene-specific “MBD scores” throughout the study. As expected based on previous studies (*3, 35*), MBD-scores were bimodally distributed (Fig. S1 A) and negatively correlated with CpG_o/e_ (Figure 1 D) providing genome-wide validation for our MBD-seq technique. The MBD-score threshold separating the two clouds of high point density in the scatter plot on Figure 1 D was used to delineate the two gene classes for computation of the “GBM class difference” (GBMcd) for each sample (Fig. 2 I, J).

### Tag-seq data processing

Transcription was quantified using Tag-seq (Meyer et al. 2011; Lohman et al. 2016). The Tag-seq reads were downloaded from the SRA database (accession SRP049522; (*23*)) and mapped against the *A.digitifera* genome using SHRiMP (*37*). Mapped reads overlapping annotated coding sequences were counted using intersection-nonempty method in HTseq version 0.6.1p1 (*38*). Normalization of raw counts and statistical analyses were performed using DESeq2 (*39*).

### Assessing variation in gene body methylation and transcription

As with gene expression analyses, normalization and statistical analyses of MBD-seq reads were performed with DESeq2 (*39*). To test for origin effects, we performed two tests that compared the two groups placed at Keppel to each other (OK vs KK) and the two groups placed at Orpheus to each other (OO vs KO). These tests were intended to identify effects of origin while controlling for environmental conditions experienced during the experiment. To assess effects of transplantation, we compared groups that originated from Orpheus to each other (OO vs OK) and groups that originated from Keppel to each other (KK vs KO). For these tests, we included an additional parameter of colony identity to identify effects of transplantation while controlling for genotype. For TagSeq, only genes with mean read count ≥3 were considered for analysis (19706 genes), and for MBD-seq we chose genes with mean read count ≥20 (27084 genes).

### Discriminant analysis of principal components

GBM data were further analyzed using discriminant analysis of principal components (DAPC) implemented in the R package adegenet (*40*), following the procedures outlined in Kenkel & Matz (2016). DAPC is a multivariate analysis method designed to identify between-group variation while neglecting within-group variation. We used this method to distill our multivariate MBD-seq dataset into single axes that maximized discrimination between natives from the two experimental sites (KK and OO samples). The result was a discriminant axis contrasting the GBM variation of the two native populations—with one pole designating native Keppel-like GBM patterns and the opposite pole designating native Orpheus-like patterns. The function was then applied to data from the transplanted samples, so that their positions along discriminant axis described their similarity in GBM patterns to the contrasted native populations. It seems plausible that, either due to natural selection or plasticity, native corals would possess patterns optimal for their particular site. If this is true, then transplants with DAPC values closer those of local corals would be expected to show improved fitness proxies (eg: samples transplanted to Keppel with DAPC values closer to the ‘Keppel-like’ pole of the discriminant axis would be expected to show higher fitness proxies and vice versa for samples transplanted to Orpheus). ‘DAPC Similarity’ values describing each transplant’s proximity along the discriminate axis to natives of their transplantation site (Fig. 4 D) were computed by taking the absolute value of the difference between the transplants’ DAPC values and the mean value for natives of their respective transplant sites, converting these distances into z-scores, and multiplied them by −1. The DAPC Similarity value for sample X was:

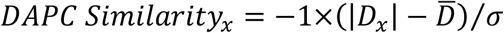

where, *D_x_* is its distance along the discriminant axis between sample X and the mean DAPC value for natives of its transplantation site, 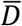 is the mean absolute distance for all transplants, and σ is the standard deviation of absolute distance for all transplants. Correlations of these values with fitness characteristics of transplanted fragments (Fig. 4 E) were computed using function *envfit* (R package *vegan* (*42*))

### Validation of MBD-seq results by bisulfite amplicon sequencing

To validate our MBD-seq results, we used targeted bisulfite sequencing. Genomic DNA for this procedure was extracted and purified as described above. It should be noted that as with the MBD-seq and Tag-seq, targeted bisulfite sequencing was performed from separate isolations from the same or closely adjacent branch. We performed bisulfite conversion using an EZ DNA Methylation-Gold kit (Zymo Research; cat. no. D5005). Thermocycler conditions for conversion were 98°C for 10 minutes followed by 53°C for 4 hours. Primers for post-conversion amplification were designed using Bisulfite Primer Seeker (Zymo Research; http://www.zymoresearch.com/tools/bisulfite-primer-seeker). Primer sequences are given in Table S1. We designed 13 primer sets to target coding sequences (excised from the *A. digitifera* reference genome) of 13 separate loci. Primer sequences included 5’ tails for downstream addition of barcodes and Illumina adapters by PCR. Target loci were selected based on either strong origin or transplant effects in the DESeq2 analyses. Amplification from bisulfite converted samples was performed using EpiMark Hot Start Taq DNA Polymerase (New England Biolabs; cat. no. M0490S). Thermocycler conditions were 95°C for 30 seconds followed by 35-40 cycles of 95°C for 15 seconds, 53°C for 30 seconds, 68°C for 30 seconds. Barcodes and adapters for multiplexed sequencing were added to the resulting PCR products in a second PCR. The oligonucleotide sequences used for barcoding were the same as those given in the Tag-seq library preparation (Meyer et al. 2011; https://github.com/z0on/tag-based_RNAseq). Sequencing was performed using paired-end 600 cycle runs on the Illumina Miseq. Resulting reads were quality trimmed using cutadapt (*30*). Quantification of CpG methylation was performed using Bismark (*43*) using a fasta file of the 13 exon sequences used to design the primers as a reference. For a given sample, CpG sites represented by fewer than 50 reads were excluded from the analysis (2022 low coverage calls excluded). The difference in means based on origin for all CpGs in each gene correlated with the difference in mean normalized counts from the MBD-seq reads (Figure 1 D).

#### Primers used for targeted bisulfite sequencing

Each primer includes a 5’ tail (bold) for amplification with barcoded primers for multiplex Illumina sequencing included in the Tag-seq library preparation.

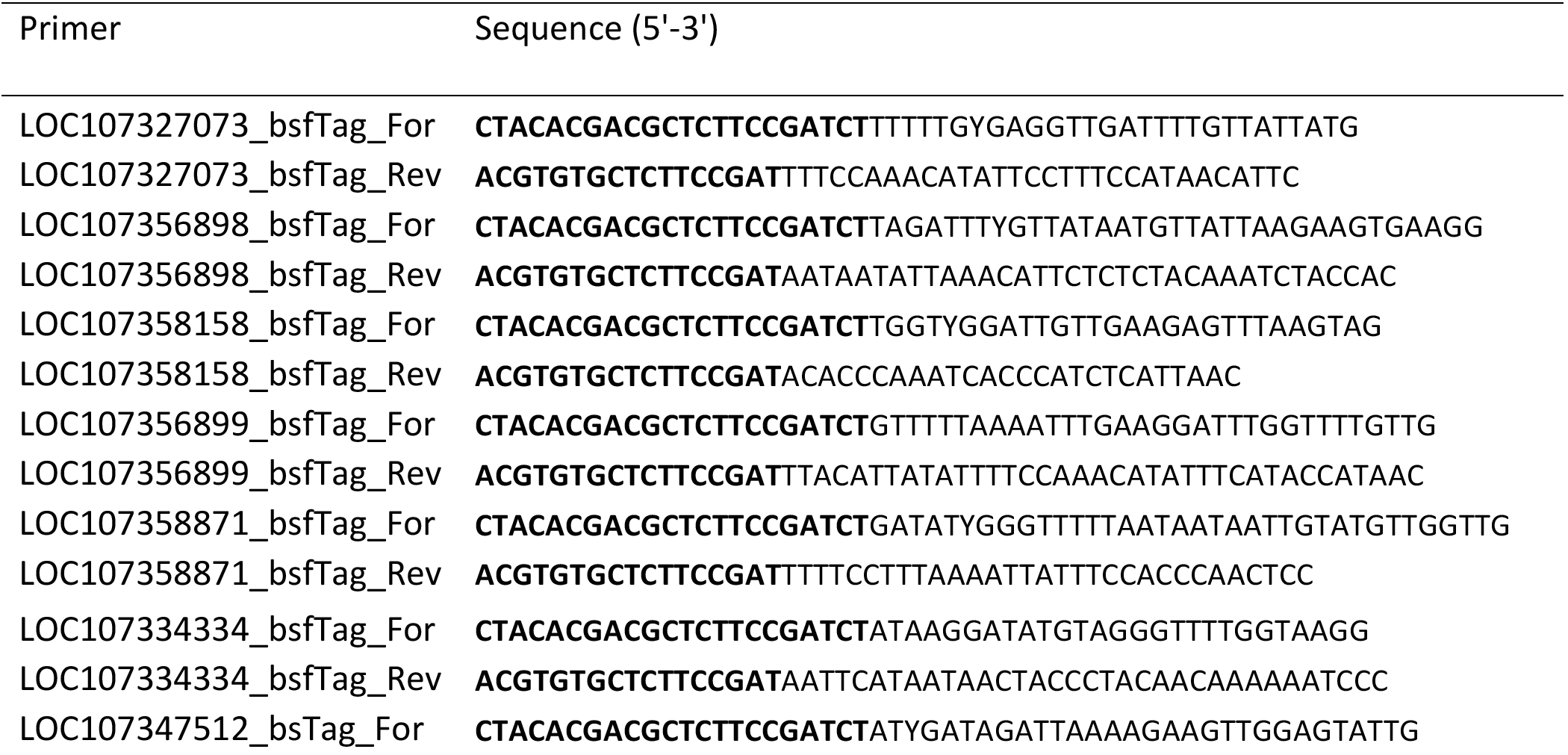
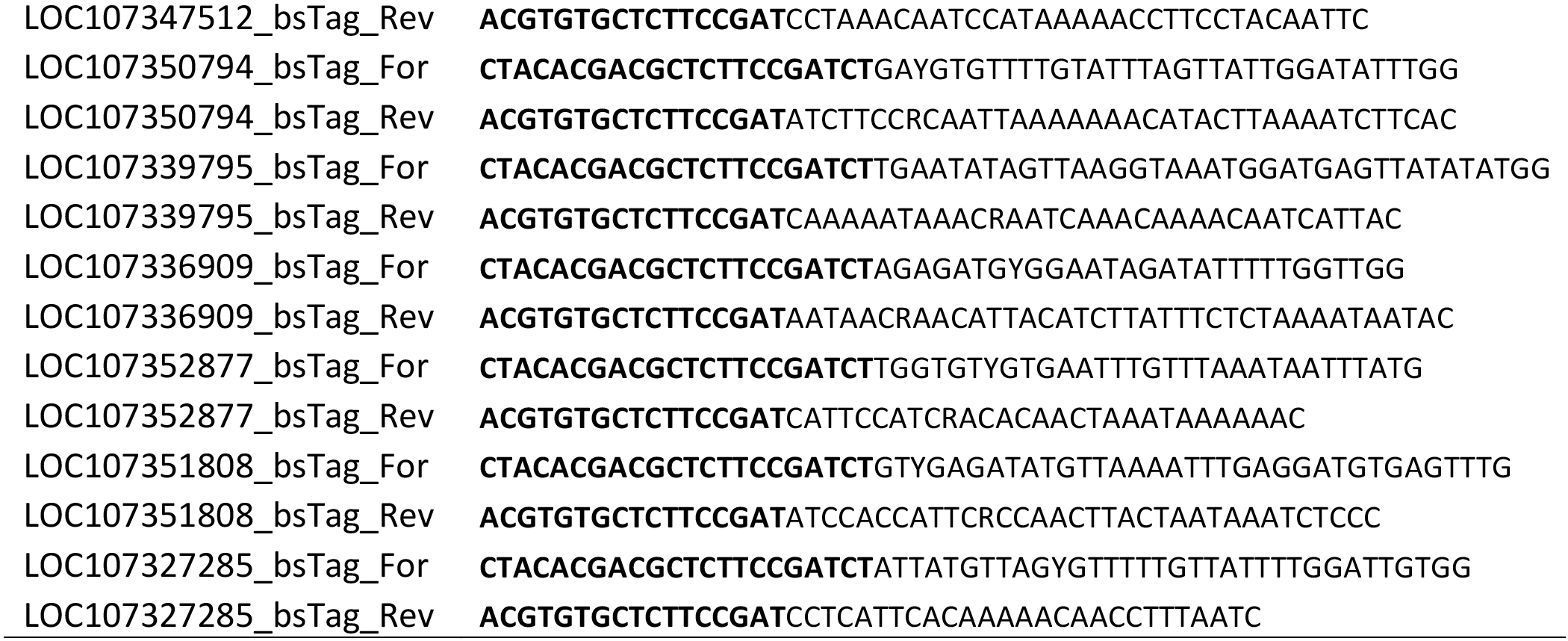

### Analysis of genetic differentiation

The MBD-seq reads were also used to analyze genetic distances among the 22 sequenced colonies. For this procedure, we concatenated reads from clone pairs into single files, and mapped the reads to the *A. digitifera* reference genome (*31*) using Bowtie2 (*32*). For the subset of 12 samples where we sequenced both captured and flow-through fractions, both sets of reads were concatenated for genotyping. Between-individual genetic distances for DAPC analysis were determined as (1 - Identity-By-State) using ANGSD (*44*) with the following filter settings: -uniqueOnly 1 -remove_bads 1 -minMapQ 20 -minQ 30 -baq 1 -minInd 10 -snp_pval 1e-2 -minMaf 0.1 (to summarize, only high-quality and uniquely-mapped reads were taken into account; only sites covered in a minimum of 10 individuals were considered; the cutoff for the p-value for the SNP being true was set at 0.01; and only alleles with frequency equal or exceeding 0.1 were used). This resulted in genotyping information for roughly 507,000 polymorphic sites. The between-population *F*_ST_ for genes passing 10% FDR cutoff for either for origin or transplant effect and a random subset of 500 non-significant genes (rather than whole genome, to speed up computation) were determined using ANGSD based on site frequency spectra as priors, which were generated for every gene subset without using filters that would affect representation of alleles of different frequencies (-uniqueOnly 1 -remove_bads 1 - minMapQ 20 -minQ 30 -baq 1 -minInd 4).

### *Determination of relative proportions of* Symbiodinium *clades*

The MBD-seq reads were mapped to the concatenated reference including A. digitifera genome and four *Symbiodinium* sp. trasncriptomes from four different genotypic groups (“clades”). Transcriptomes for *Symbiodinium* clades A and B were from (*45*) and transcriptomes for clades C and D were from (*46*). We then counted the relative proportions of reads producing highly unique matches (mapping quality 40 or higher) to each *Symbiodinium* transcriptome, using a custom perl script *zooxType.pl*. All corals were found to be dominated by *Symbiodinium* clade C.

### Data and scripts availability

TagSeq, MBD-seq, and amplicon-bisulfite sequencing datasets are available through NCBI Short Read Archive (Project Accession: SRP049522). Scripts and traits data are available in a GitHub repository dedicated to this paper (https://github.com/grovesdixon/reciprocal_transplant_methylation).

## Supplementary Tables and Figures

**Table S1.** Sample sizes

| Sample group | N (TagSeq) | N (MBD-seq, captured and flow-through fractions) | N (MBD-seq, only captured fraction) |
| --- | --- | --- | --- |
| KK | 14 | 3 | 8 |
| KO | 15 | 3 | 8 |
| OK | 15 | 3 | 8 |
| OO | 15 | 3 | 8 |

**Figure S1.**
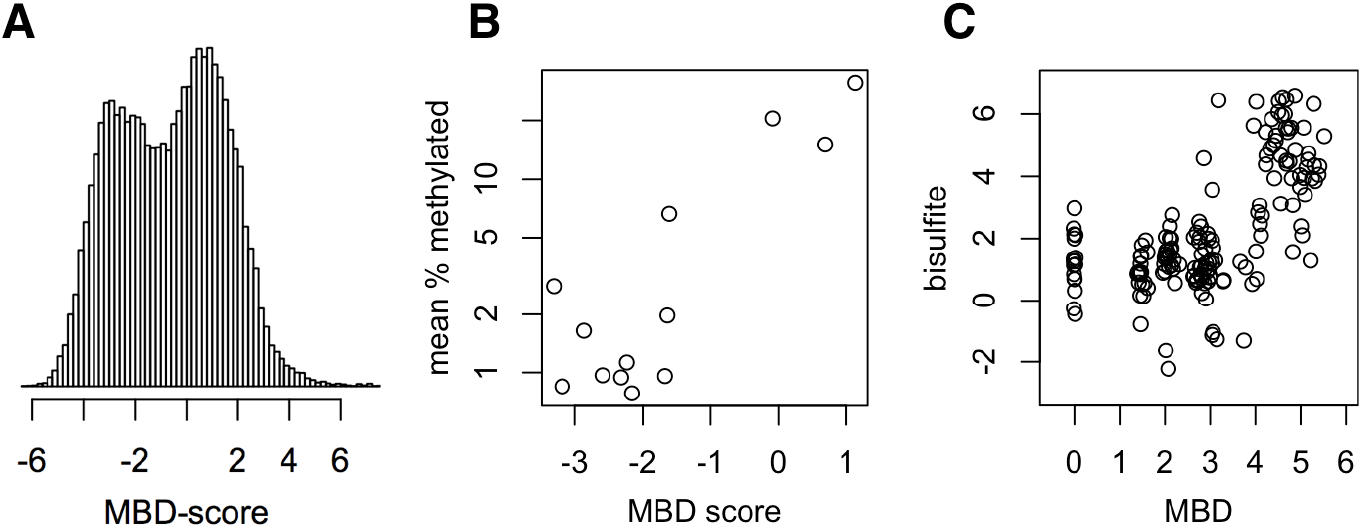
Distribution of MBD-scores (A) and validation by bisulfite sequencing of amplicons (B, C). B: Mean percent methylation was calculated as the proportion of methylated CpG sites within each gene averaged across all samples. C: Correlation between MBD-based and bisulfite-based measurement of the effects of each coral individual on methylation for each of the assayed genes.

**Figure S2.**
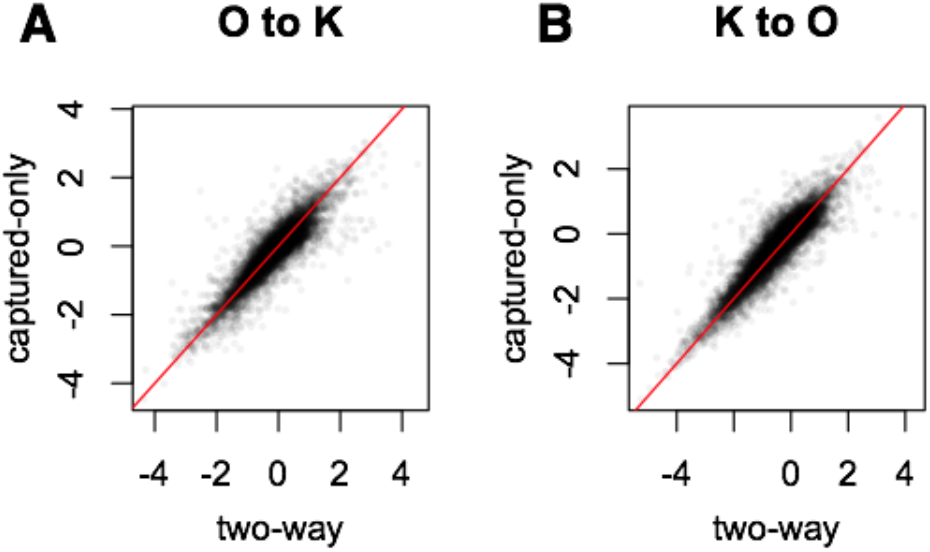
Correlation between inferred changes in GBM based on comparison of MBD-captured and flow-through libraries (x-axis) and only on MBD-captured fraction (y-axis). The red line is 1:1 correspondence. (A) Orpheus to Keppel transplantation; (B) Keppel to Orpheus transplantation. These plots were generated for a subset of 12 samples (three for each of the four experimental groups, KK, KO, OK, and OO) for which both the captured and the flow-through fractions were sequenced.

**Figure S3.**
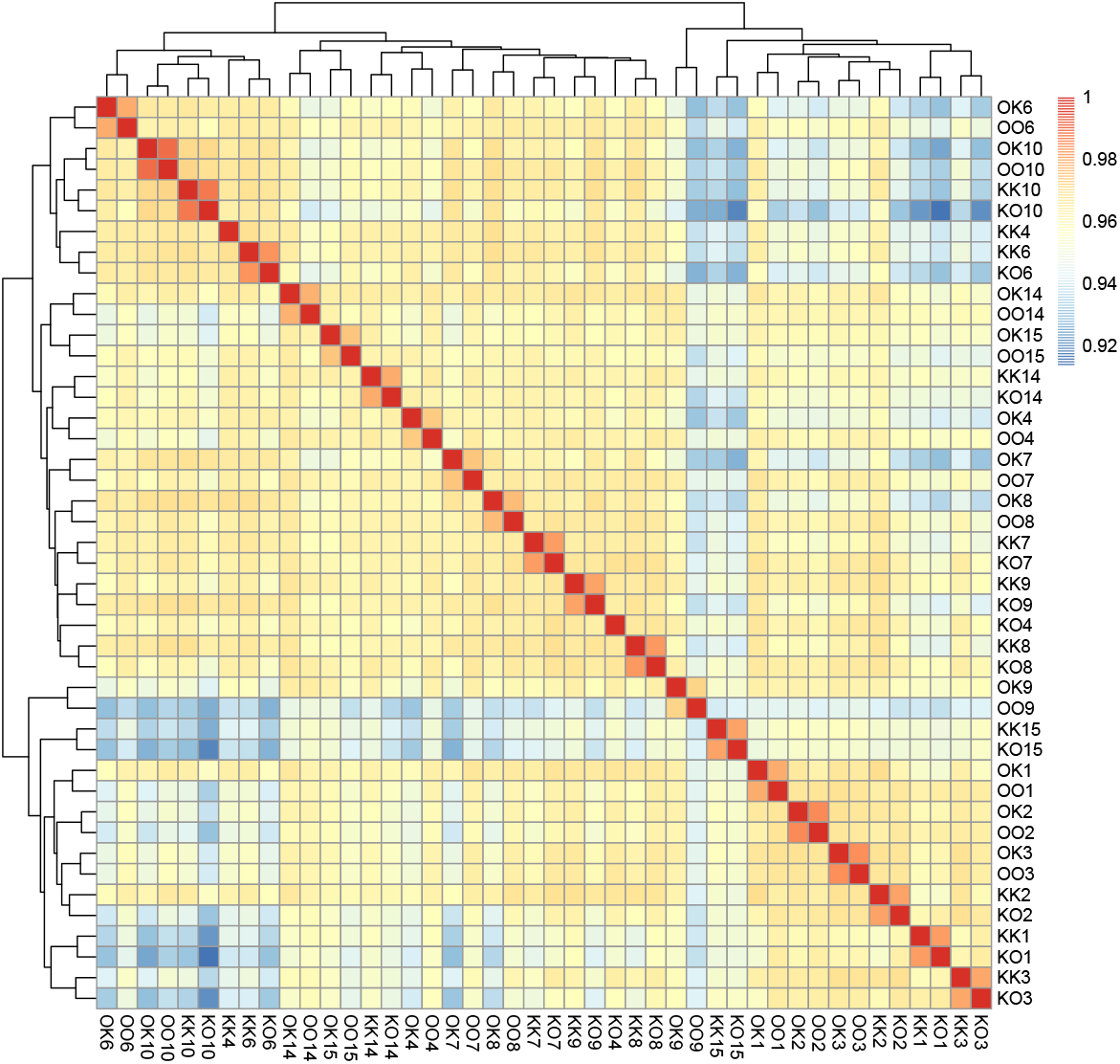
Heatmap of GBM correlations among samples, illustrating strong dependence of GBM on genotype. Colors indicate Spearman’s rank correlations for normalized MBD-seq read counts across all coding genes (N = 24853); the lowest observed correlation value was 0.9. Samples were clustered by maximum distance method. First letter of sample names indicates sample origin, second letter indicates transplantation site, and number indicates replicate. Samples sharing the same first letter and the same number are clonal fragments from the same colony. All of the 22 clone pairs except one (sample pair KK4 and KO4) clustered together.

**Figure S4.**
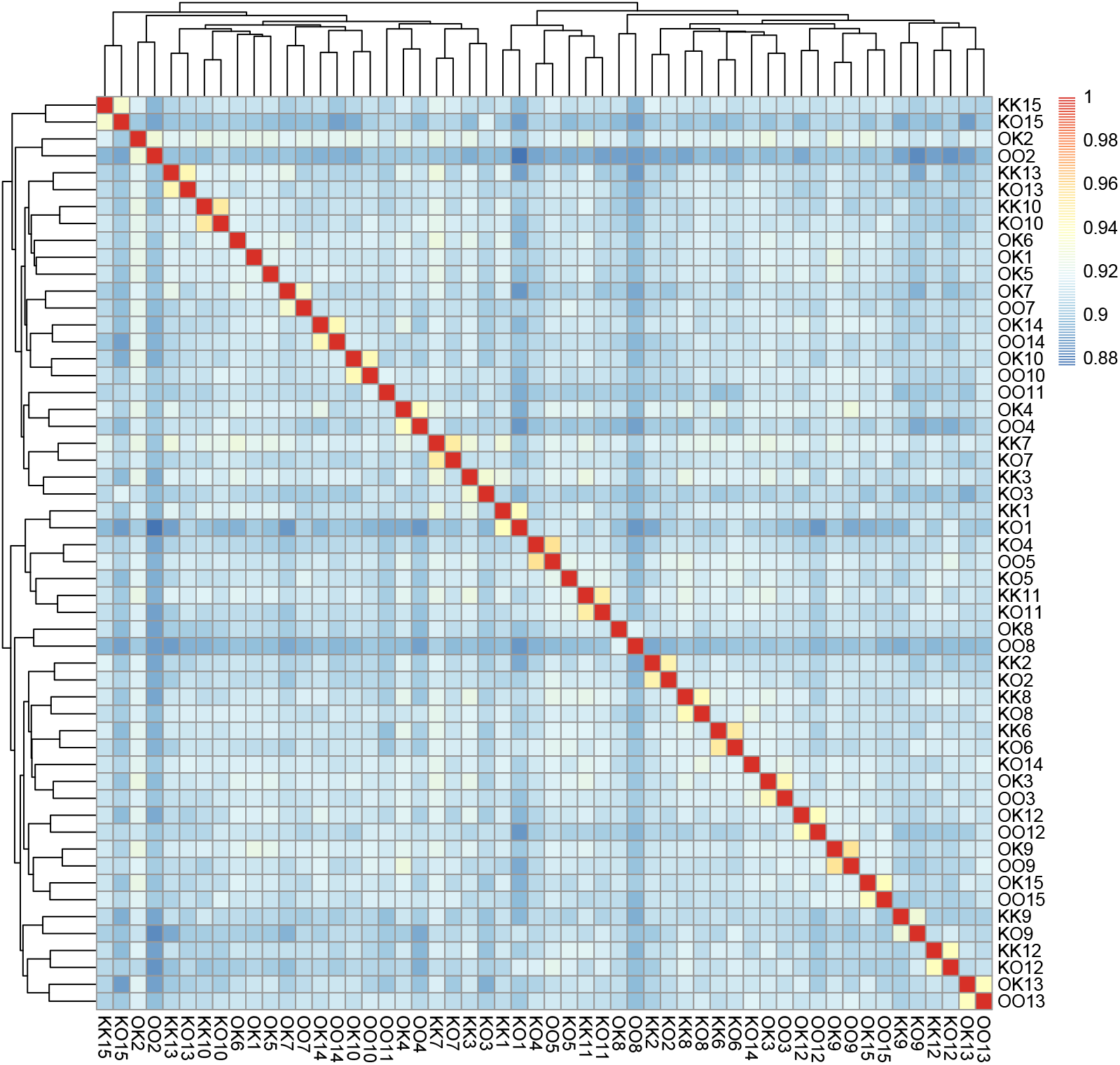
Heatmap of overall correlations in transcription illustrating strong dependence of transcription on genotype. Colors indicate Spearman’s correlations for normalized Tag-seq read counts across all coding genes (N=19706). Samples were clustered by maximum distance method. First letter of sample names indicates sample origin, second letter indicates transplantation site, and number indicates replicate. Samples sharing the same first letter and number are clonal fragments from the same colony. All 24 available clone pairs clustered together (six samples lacked data for clone pairs).

**Figure S5.**
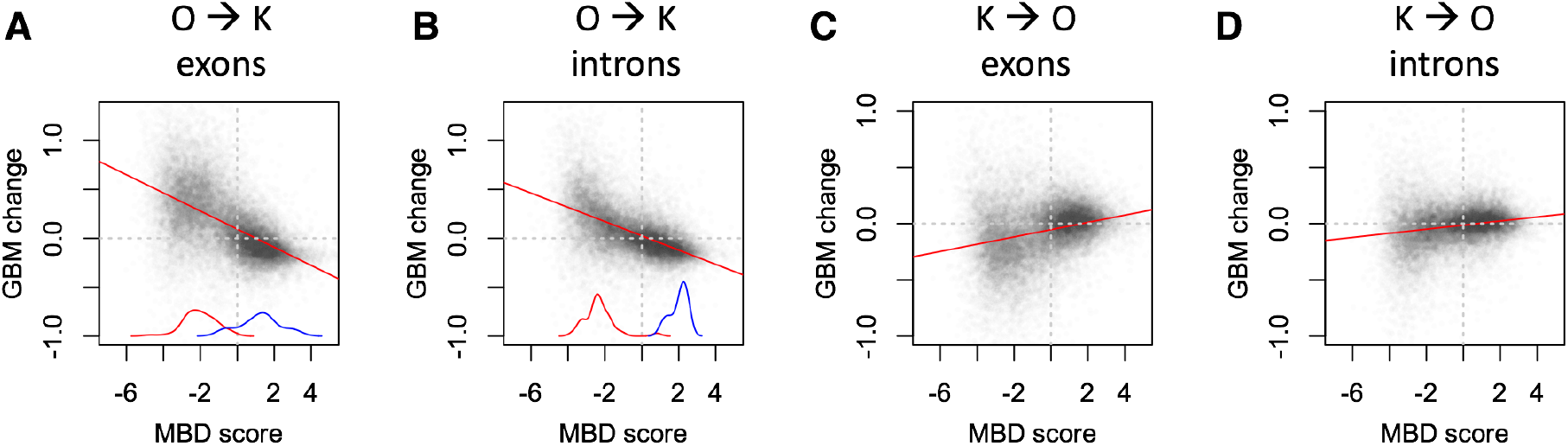
GBM log_2_ fold-changes change in response to transplantation in exons and introns plotted against MBD-score for each gene. Red and blue curves near the x-axis are density plots of significantly up- and down-regulated genes.

**Figure S6.**
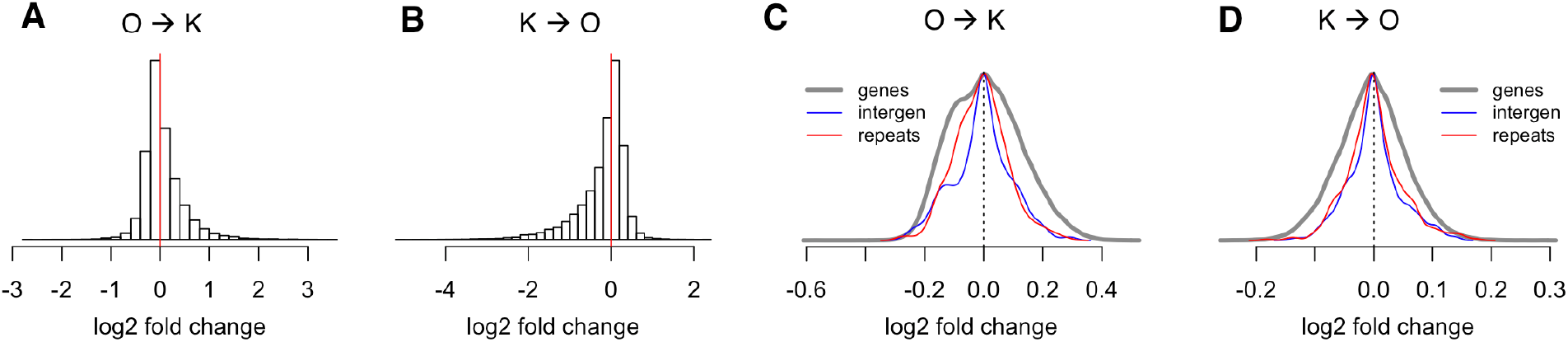
Absolute methylation changes in response to transplantation (“away” vs. “home”) can be positive or negative across genes (A, B) and affect mostly protein-coding regions (C, D). Plots on A and B are based on comparison between captured and flow-through fractions for 12 samples (three for each of the four experimental groups, KK, KO, OK, and OO). Plots on panels C and D were generated based on captured-only analysis of all 44 MBD-seq samples (Table S1).

**Figure S7.**
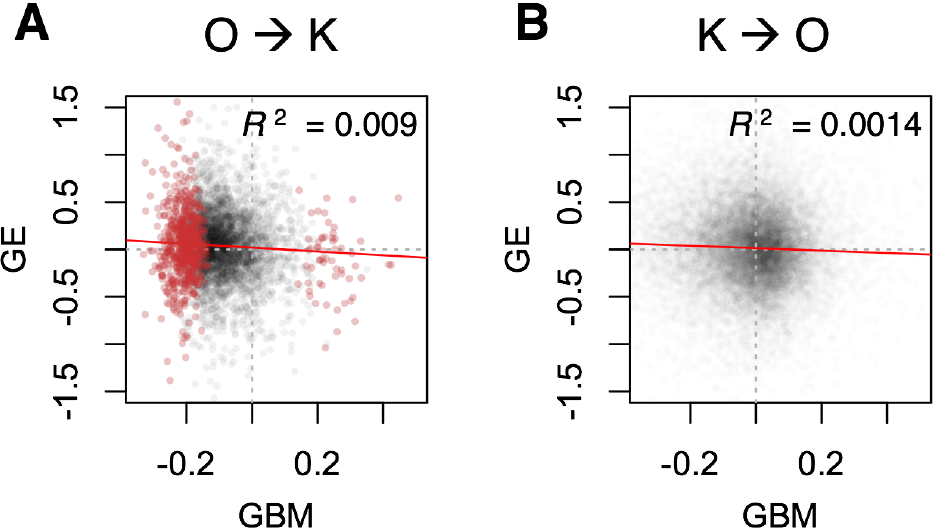
Gene expression changes in response to transplantation are negatively correlated with GBM changes. Both correlations are highly significant (p<0.0001). Red points on panel A represent genes with statistically significant (FDR<10%) change in GBM.

**Figure S8.**
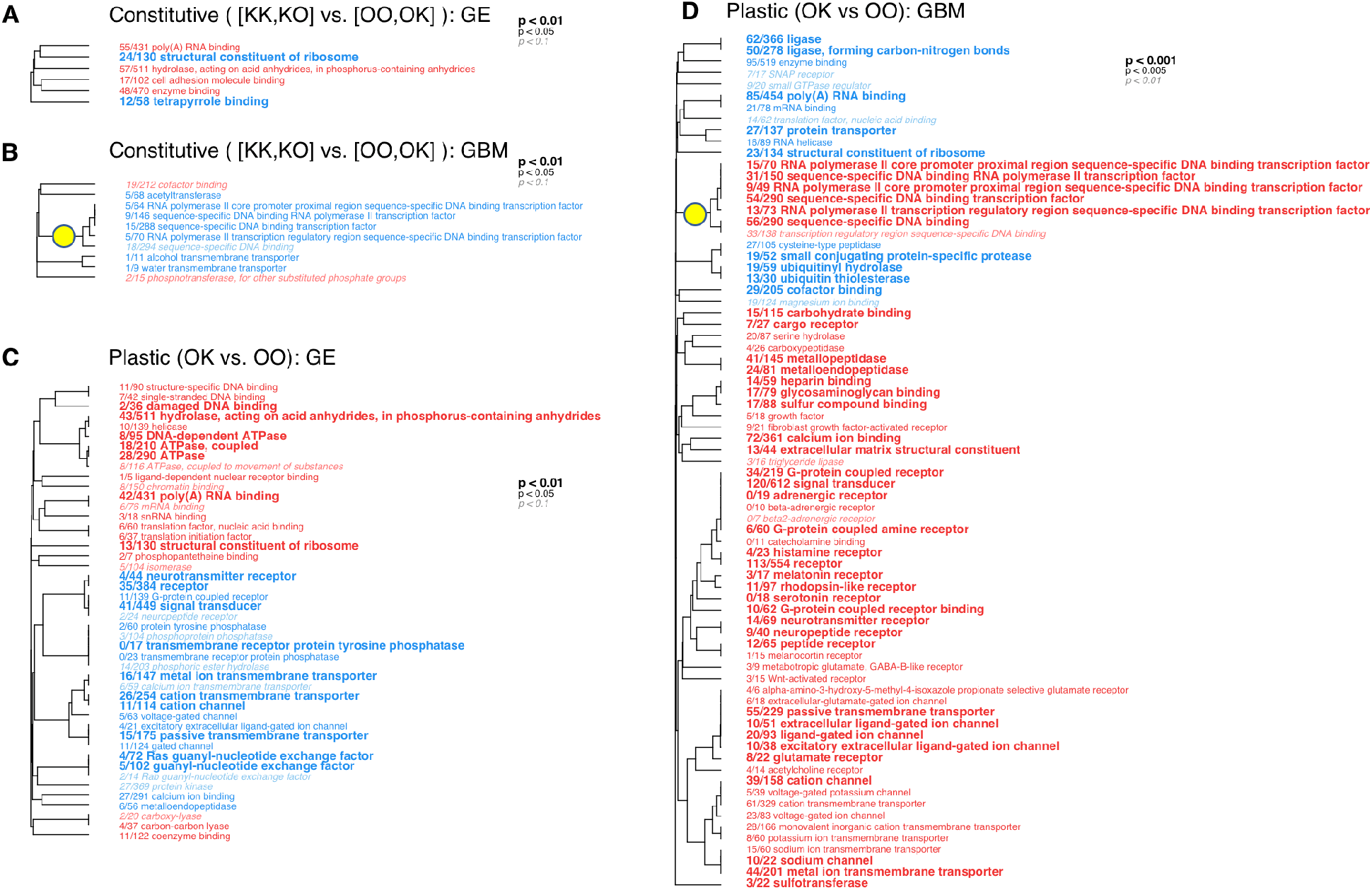
Gene ontology categories significantly enriched with up- or down-regulated genes (red and blue, respectively) in GE and GBM comparisons. The Benjamini-Hochberg corrected significance of enrichment is indicated by the size and type of the font (see legends). Note that for plastic GBM the significance thresholds are ten times more stringent, to fit the GO plot onto one page. The fraction preceding the category name is the number of genes exceeding raw p-value of 0.05 relative to the total number of genes in the category. The dendrograms are hierarchical clustering of GO categories based on proportion of shared genes (*47*). Genes comprising the branch labelled by the yellow circle (transcription factors) show plastic changes (D) that are opposite to the constitutive differences between populations (B).

**Figure S9.**
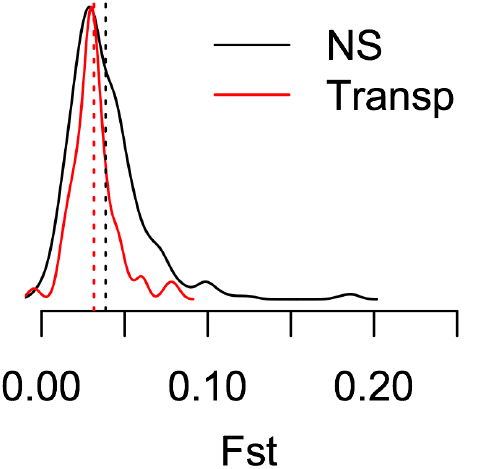
Density plots of *F*_ST_ for genes that show significant (FDR <0.1) GBM change in response to transplantation from Orpheus to Keppel (“Transp”, red curve) compared to 500 randomly picked non-significant genes (“NS”, black curve). Dashed vertical lines mark the mean of each dataset.

